# Effects of *Cyperus Esculentus* Infused Diet on the Hippocampus in Rotenone-Induced Parkinson-like Disease Model in adult male Wistar Rats; a preliminary study

**DOI:** 10.64898/2026.10.13.758640

**Authors:** Dada Moyosoluwa Mary, Adelodun Stephen Taiye, Awolusi Oluwabukunmi Eniola

**Affiliations:** Department of Anatomy, School of Basic Medical Sciences, Benjamin Carson Snr College of Health and Medical Sciences, Babcock University Ilishan-Remo, Ogun State, Nigeria

**Keywords:** Parkinson’s disease, rotenone, *Cyperus esculentus*, hippocampus, neuroprotection, neurodegeneration

## Abstract

Parkinson’s disease is a progressive neurodegenerative disorder contributing to cognitive decline and memory impairment beyond its characteristic motor deficits. Rotenone, a mitochondrial complex I inhibitor, reliably reproduces both the motor and non-motor features of Parkinson’s disease in experimental models. Tiger nut (*Cyperus esculentus*) is a commonly consumed plant known for its antioxidant and anti-inflammatory properties; however, its neuroprotective efficacy against rotenone-induced hippocampal damage is not yet fully elucidated. This study, therefore, evaluated the neuroprotective efficacy of tiger nut-infused diet on the hippocampus in a rotenone-induced Parkinson’s disease model using male Wistar rats. Thirty-two male Wistar rats weighing between 100 g to 150 g were obtained from the Animal Holding facility of Babcock University, Ilishan-Remo, Ogun State. The rats were randomly assigned into four groups (A, B, C and D) (n = 8). Group A served as the control and received normal rat chow and sunflower oil. Group B received tiger nut-infused diet only. Group C received rotenone (2 mg/kg) intraperitoneally. Group D received rotenone (2 mg/kg) concurrently with tiger nut-infused diet for 28 days. Neurobehavioral assessments were conducted using the open field test and Y-maze tests. Oxidative stress biomarkers, including malondialdehyde, catalase, and glutathione, were assayed from hippocampal homogenates, and gamma-aminobutyric acid was measured as a neurotransmitter parameter. Histological evaluation was performed using hematoxylin and eosin for general cytoarchitecture, Cresyl Fast Violet for Nissl substance integrity, and Luxol Fast Blue for myelin visualization. Data were analyzed using One-way ANOVA followed by the Student’s Newman-Keuls post-hoc test on GraphPad Prism 9.0, with significance set at p ≤ 0.05. Rotenone administration produced significant impairments in locomotor activity and spatial working memory, reduced catalase activity, and caused marked histopathological alterations in the hippocampus, including neuronal vacuolation, chromatolysis, and myelin disruption. Tiger nut-infused diet partially attenuated these effects, most notably by enhancing catalase activity and preserving hippocampal cytoarchitecture compared to the rotenone-only group. This study concludes that tiger nut-infused diet demonstrates partial neuroprotective potential against rotenone-induced hippocampal neurotoxicity, with implications for dietary intervention in Parkinson-like disease-related cognitive decline. Further studies involving phytochemical screening of the tiger nut-infused diet are recommended to identify and isolate the specific active ingredient responsible for its neuroprotective effects, enabling investigation of its mechanisms of action at the molecular level.

## 1. Introduction

Parkinson’s disease (PD) is a progressive neurodegenerative disorder characterized primarily by the loss of dopaminergic neurons in the substantia nigra, resulting in classical motor symptoms including resting tremor, rigidity, bradykinesia, and postural instability [1]. Beyond these motor manifestations, PD is increasingly recognized as a disease with significant non-motor burden, including cognitive decline, memory impairment, and emotional disturbances, many of which have been linked to pathological involvement of limbic structures, particularly the hippocampus [2]. As the second most prevalent neurodegenerative disorder worldwide, PD affects over 10 million individuals globally, with its incidence rising sharply in aging populations and projections suggesting a doubling of its global burden by 2040 [3, 4, 5].

The hippocampus, a bilateral structure located in the medial temporal lobe, plays a central role in learning, memory consolidation, and spatial navigation [6]. In Parkinson’s disease, the progressive spread of Lewy body pathology from subcortical regions to limbic and cortical structures directly implicates the hippocampus in the cognitive and neuropsychiatric features of the disease [7]. Rotenone-induced neuroinflammation has been shown to impair synaptic plasticity, reduce neurogenesis in the dentate gyrus, and disrupt hippocampal neuronal integrity, collectively contributing to the memory and learning deficits observed in experimental PD models [8]. These findings underscore the hippocampus as a critical, yet often underemphasized, target of PD-related neurodegeneration.

Rotenone, a naturally occurring pesticide and mitochondrial complex I inhibitor, is widely employed as an experimental model of PD owing to its ability to reliably reproduce both the motor and non-motor features of the disease [9]. Its mechanism involves the generation of reactive oxygen species, induction of oxidative stress, activation of neuroinflammatory cascades, and progressive dopaminergic neuronal death, closely mirroring the neuropathological processes observed in human PD [10]. Epidemiological studies have further associated occupational rotenone exposure with an elevated risk of developing Parkinson’s disease in humans, strengthening the translational validity of this experimental model [11].

Current pharmacological management of PD, centered primarily on dopamine replacement therapy with levodopa, provides symptomatic relief for motor deficits but does not halt the underlying neurodegenerative process and offers limited efficacy against hippocampus-related cognitive impairment [12]. This therapeutic gap has fueled growing interest in natural, plant-derived compounds with neuroprotective potential. Tiger nut (*Cyperus esculentus*), a perennial sedge belonging to the family Cyperaceae and widely cultivated and consumed across West Africa and the Mediterranean region, has attracted considerable scientific attention for its rich array of bioactive phytochemicals, including polyphenols, flavonoids, alkaloids, saponins, and tannins [13, 14]. These constituents have been shown to possess antioxidant, anti-inflammatory, and neuroprotective properties that may mitigate oxidative neuronal damage and support cognitive function [15, 16].

Despite its promising pharmacological profile, the specific neuroprotective efficacy of a tiger nut-infused diet against rotenone-induced hippocampal damage remains insufficiently characterized. Therefore, the present study investigated the effects of tiger nut (*Cyperus esculentus*)-infused diet on the hippocampus in a rotenone-induced Parkinson’s disease model using male Wistar rats, by evaluating neurobehavioral performance, hippocampal oxidative stress biomarkers, neurotransmitter levels, histomorphological integrity, and parvalbumin immunoexpression.

## 2. Materials and Methods

### 2.1 Ethical approval

Ethical clearance for this research was granted by the Babcock University Health Research Ethical Committee (BUHREC) under the protocol reference (BUHREC 0323/26), prior to commencement of all experimental procedures. All animal handling and experimental protocols were conducted in strict accordance with international standards for welfare and the institutional policies of Babcock University.

### 2.2 Preparation of Tiger Nut-Infused Diet

Tiger nut (Cyperus esculentus) tubers were obtained from Ilishan Market, Ogun State, Nigeria. The tubers were thoroughly washed, air-dried, weighed, and incorporated into the feed formulation at 11% by weight, replacing a proportional quantity of corn oil and other constituents, following the modified method [17]. The control diet was prepared identically but without the tiger nut component.

TABLE SHOWING THE COMPOSITION OF NORMAL RAT CHOW AND CYPERUS ESCULENTUS INFUSED CHOW.

| INGREDIENTS | % OF WEIGHT | CONTROL DIET<br>AMOUNT/100g | TIGER NUT DIET<br>AMOUNT/100g |
| --- | --- | --- | --- |
| Groundnut cake | 25 | 25 | 24.4 |
| Palm kernel cake | 25 | 25 | 24.4 |
| Corn Starch | 15 | 15 | 13.5 |
| Industrial Soya | 10 | 10 | 9.4 |
| Bone meal | 5 | 5 | 5 |
| Wheat (fibre) | 5 | 5 | 4.8 |
| Lysine | 0.2 | 0.2 | 0.2 |
| Methionine | 0.3 | 0.3 | 0.3 |
| Concentrate | 4.5 | 4.5 | 4.5 |
| Ground tiger nut |  | 0 | 11 |
| Corn oil | 10 | 10 | 2.63 |
| Total | 100 | 100 | 100.1 |

Each cage received a pre-weighed amount of tiger nut-infused chow daily. After 24 hrs, the remaining feed was weighed to determine the total consumption. Although, individual feed intake may not be accurately determined, an average consumption was estimated by dividing the total intake by the number of rats per cage.

### 2.3 Experimental Animals

Thirty-two male Wistar rats weighing between 100 and 150 g were obtained from the Animal Holding Facility of Babcock University, Ilishan-Remo, Ogun State, Nigeria. The animals were housed in well-ventilated plastic cages under standard laboratory conditions and allowed to acclimatize for seven days prior to the commencement of the experiment. Cage bedding, consisting of wood shavings, was replaced every two days to prevent ammonia accumulation and maintain aseptic conditions. Standard rat chow and clean water were provided twice daily throughout the acclimatization and experimental periods.

### 2.4 Experimental Design and Treatment

Following acclimatization, the thirty-two rats were randomly assigned into four groups of eight animals each (n = 8) and treated for 28 days as follows:

• Group A (Control): received normal rat chow and 0.5 ml of sunflower oil intraperitoneally daily.

• Group B (Tiger nut only): received tiger nut-infused diet for the duration of the experiment [18].

• Group C (Rotenone only): received intraperitoneal administration of rotenone at 2 mg/kg/day [19].

• Group D (Rotenone plus tiger nut): received intraperitoneal rotenone at 2 mg/kg/day concurrently with tiger nut-infused diet for 28 days [18, 19].

Body weights were recorded weekly throughout the experiment using a weighing balance, with the initial weight measured during acclimatization and the final weight recorded on the day prior to sacrifice.

### 2.5 Neurobehavioral Studies

Neurobehavioral assessments were conducted prior to sacrifice using the Open Field Test and the Y-Maze Test to evaluate locomotor activity, exploratory behavior, and spatial working memory.

The Open Field Test (OFT) was used to assess exploratory behavior, general motor activity, and anxiety-related responses. The apparatus consisted of a square enclosure measuring 72 × 72 cm with 36 cm surrounding walls. The floor was divided into sixteen 18 × 18 cm squares using red lines, with a central square demarcated using a blue marker. One wall of the apparatus was constructed from plexiglass to permit visual observation. A video camera was mounted above the arena and connected to tracking software under uniform illumination. Each rat was placed in the center of the apparatus and allowed to explore freely for five minutes while behavior was recorded. Parameters assessed included locomotor activity, rearing, grooming, freezing, center square entries, defecation, and urination. The apparatus was thoroughly cleaned with 70% ethanol between trials to eliminate olfactory cues.

Spatial working memory was evaluated using the Y-Maze Test, which exploits the natural tendency of rodents to explore novel environments [25]. The apparatus consisted of three identical arms arranged symmetrically at 120° from one another, each approximately 30–40 cm in length, 8–10 cm in width, and 12–15 cm in height. Each rat was placed at the end of one arm and allowed to explore the maze freely for five minutes while its movements were recorded by an overhead camera connected to tracking software. An arm entry was defined as the placement of all four limbs within an arm. Spontaneous alternation, used as an index of spatial working memory, was calculated as the ratio of actual alternations (consecutive entries into all three arms without repetition) to the maximum possible alternations, expressed as a percentage. Total arm entries were also recorded as a measure of locomotor activity. The maze was wiped with 70% ethanol between trials to remove olfactory cues.

### 2.6 Sacrifice and Tissue Harvest

At the conclusion of the experimental period, all animals were euthanized by cervical dislocation following completion of neurobehavioral assessments. The whole brain was immediately excised and weighed. Brain tissue allocated for biochemical and neurotransmitter assays was placed in phosphate buffer saline (PBS), while tissue for histological and immunohistochemical analysis was fixed in 10% neutral buffered formalin. Animal carcasses were disposed of by burial in a designated area adjacent to the animal facility in accordance with institutional biosafety guidelines.

### 2.7 Histological Processing and Staining

Brain tissue designated for histological examination was processed through standard paraffin embedding procedures. Briefly, tissues were fixed in 10% neutral buffered formalin, dehydrated through ascending concentrations of ethanol (70%, 90%, and 100%), cleared in three changes of xylene (20 minutes each), and infiltrated with molten paraffin wax at 60°C. Tissues were then embedded in paraffin blocks, sectioned at 30 µm for histology and 15 µm for immunohistochemistry using a rotary microtome (Leica RM2245), and mounted on glass slides (Thermo-Scientific). Three staining techniques were applied to hippocampal sections: Hematoxylin and Eosin (H&E) for general cytoarchitecture, Luxol Fast Blue (LFB) for myelin visualization, and Cresyl Fast Violet (CFV) for the demonstration of Nissl substance integrity.

For H&E staining, sections were deparaffinized in xylene, rehydrated through graded alcohols, overstained with Harris hematoxylin for 5–7 minutes, differentiated in acid alcohol and Scott’s water, counterstained with 1% eosin for 10 minutes, dehydrated, and cleared in xylene prior to mounting.

For LFB staining, sections were deparaffinized, rehydrated through 100% and 95% ethanol, and incubated in 0.1% Luxol Fast Blue solution for 2 hours at 60°C. Sections were then differentiated individually in 0.05% lithium carbonate solution and 70% ethanol, counterstained with Cresyl Fast Violet for 8 minutes, dehydrated, cleared in xylene, and mounted with DPX.

For CFV staining, tissue sections were mounted on gelatin-coated slides, oven-dried overnight, and subjected to a de-fat step in 1:1 alcohol/chloroform overnight before rehydration. Sections were stained in warm 0.1% Cresyl Violet solution at 37°C for 7 minutes, differentiated in 95% ethanol for 20 minutes, dehydrated in 100% alcohol, cleared in xylene for 5 minutes, and examined under the microscope.

### 2.8 Biochemical Assay

The hippocampus was dissected from each brain, placed in normal saline, and homogenized to obtain a uniform tissue homogenate. The homogenate was centrifuged at 3,000 rpm for 20 minutes using a Model 90-1 centrifuge, and the resulting supernatant was carefully collected and stored at −4°C until analysis. The following oxidative stress biomarkers were assessed spectrophotometrically.

Malondialdehyde (MDA) levels were determined by the thiobarbituric acid reactive substances method. Briefly, 100 µL of sample was combined with 500 µL each of phosphate-buffered saline and trichloroacetic acid, incubated at room temperature, and centrifuged for 90 seconds at 4,000 rpm. One millilitre of 0.33% thiobarbituric acid was added, and the mixture was heated in a boiling water bath for one hour. After cooling, absorbance was measured at 533 nm.

Catalase (CAT) activity was measured by combining 100 µL of sample with 500 µL each of phosphate-buffered saline and 0.03% hydrogen peroxide. After a 10-minute incubation at room temperature, the reaction was terminated with 500 µL of 1% potassium dichromate in acetic acid. The mixture was heated in a boiling water bath for 10 minutes, cooled, and absorbance measured at 570 nm.

Glutathione (GSH) concentration was determined by combining 100 µL of sample with 500 µL each of phosphate-buffered saline and 5,5’-dithiobis-(2-nitrobenzoic acid) (DTNB) solution. After 15 minutes of incubation at room temperature, the resulting yellow complex was measured at 412 nm.

### 2.9 Neurotransmitter Assay

Gamma-aminobutyric acid (GABA) levels in hippocampal tissue were assessed using a commercial GABA detection kit. Tissue samples were washed in cold phosphate-buffered saline, resuspended in 100 µL of GABA Assay Buffer, homogenized, and incubated on ice for 15–30 minutes before centrifugation at 4°C at maximum speed. The resulting supernatant was collected, and a 100 µL reaction mixture containing GABA enzyme mix, developer, and probe was added to each sample well per the kit protocol. Samples were incubated at 37°C for 30–60 minutes in the dark, and absorbance was measured at 450 nm. GABA concentrations were calculated from a standard curve prepared alongside the samples.

### 2.10 Immunohistochemical Assay

Parvalbumin (PV) immunoexpression was evaluated in hippocampal sections to assess the integrity of GABAergic inhibitory interneurons [36]. Brain tissues were fixed in 4% paraformaldehyde at 4°C for 24–48 hours, processed in paraffin, and sectioned at 5–10 µm. Sections were deparaffinized in xylene, rehydrated through graded ethanol to distilled water, and subjected to antigen retrieval in citrate buffer (pH 6.0) by heating for 10–20 minutes. Non-specific binding was blocked with 5% normal serum or bovine serum albumin (BSA) in PBS for 30–60 minutes at room temperature. Sections were then incubated overnight at 4°C with anti-Parvalbumin primary antibody diluted in blocking buffer, washed three times in PBS, and incubated with horseradish peroxidase (HRP)-conjugated secondary antibody for one hour at room temperature. Following further PBS washes, DAB chromogen was applied to visualize Parvalbumin-positive sites, producing a brown precipitate. Sections were counterstained with hematoxylin, dehydrated, cleared in xylene, and cover slipped with permanent mounting medium. Negative controls were processed identically but without primary antibody.

### 2.11 Photomicrography and Image Analysis

Stained slides were examined under a Leica DM750 light microscope equipped with a Leica ICC50 digital camera. Photomicrographs were acquired and imported into ImageJ software for histomorphometric analysis and quantification of staining intensity and cellular density.

### 2.12 Statistical Analysis

All data were expressed as mean ± standard error of the mean (SEM) and analyzed using one-way analysis of variance (ANOVA) followed by the Student’s Newman-Keuls post-hoc multiple comparisons test on GraphPad Prism version 9.0. Statistical significance was set at p ≤ 0.05.

## 3. Results

### 3.1 Morphological Analysis

Figure 1a and 1b show the change in body weight and brain weight of the rats across the experimental groups respectively. The results showed a significant increase in body weight of the control group when compared with the tiger nut only, rotenone only, and rotenone plus tiger nut groups. The rotenone plus tiger nut group showed a significant reduction in body weight when compared with both the rotenone only and tiger nut only groups. Regarding brain weight, the results showed a significant increase in the control group when compared with the tiger nut only group, while both the rotenone only and rotenone plus tiger nut groups showed significantly higher brain weights compared to the control. The tiger nut only group recorded the lowest brain weight, showing a significant decrease when compared with all other groups.

**Figure 1.**
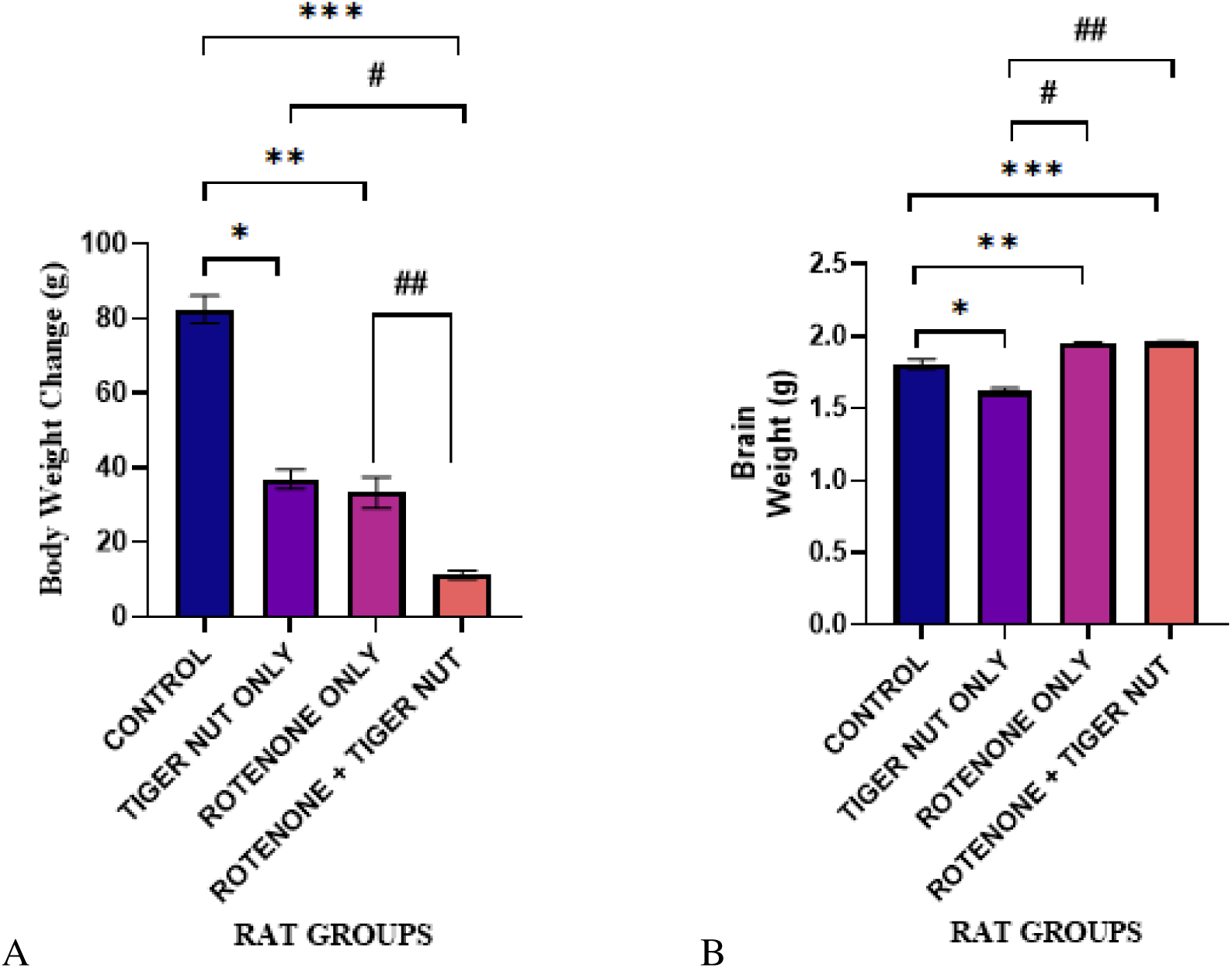
(a & b): Bar charts showing (a) the change in body weight and (b) the brain weight (g) of the rats across experimental groups. Values are presented as Mean ± SEM. P ≤ 0.05.

### 3.2 Neurobehavioral Assessment

#### 3.2.1 Y-Maze Test

Figure 2a and 2b show the total arm entries and percentage of alternations of the rats across the experimental groups respectively. The results showed a significant increase in total arm entries in the control and tiger nut only groups when compared with the rotenone only and rotenone plus tiger nut groups. Similarly, the results showed a significant increase in the percentage of alternations in the control and tiger nut only groups when compared with the rotenone only and rotenone plus tiger nut groups. The rotenone plus tiger nut group did not show significant recovery compared to the rotenone only group across both parameters.

**Figure 2.**
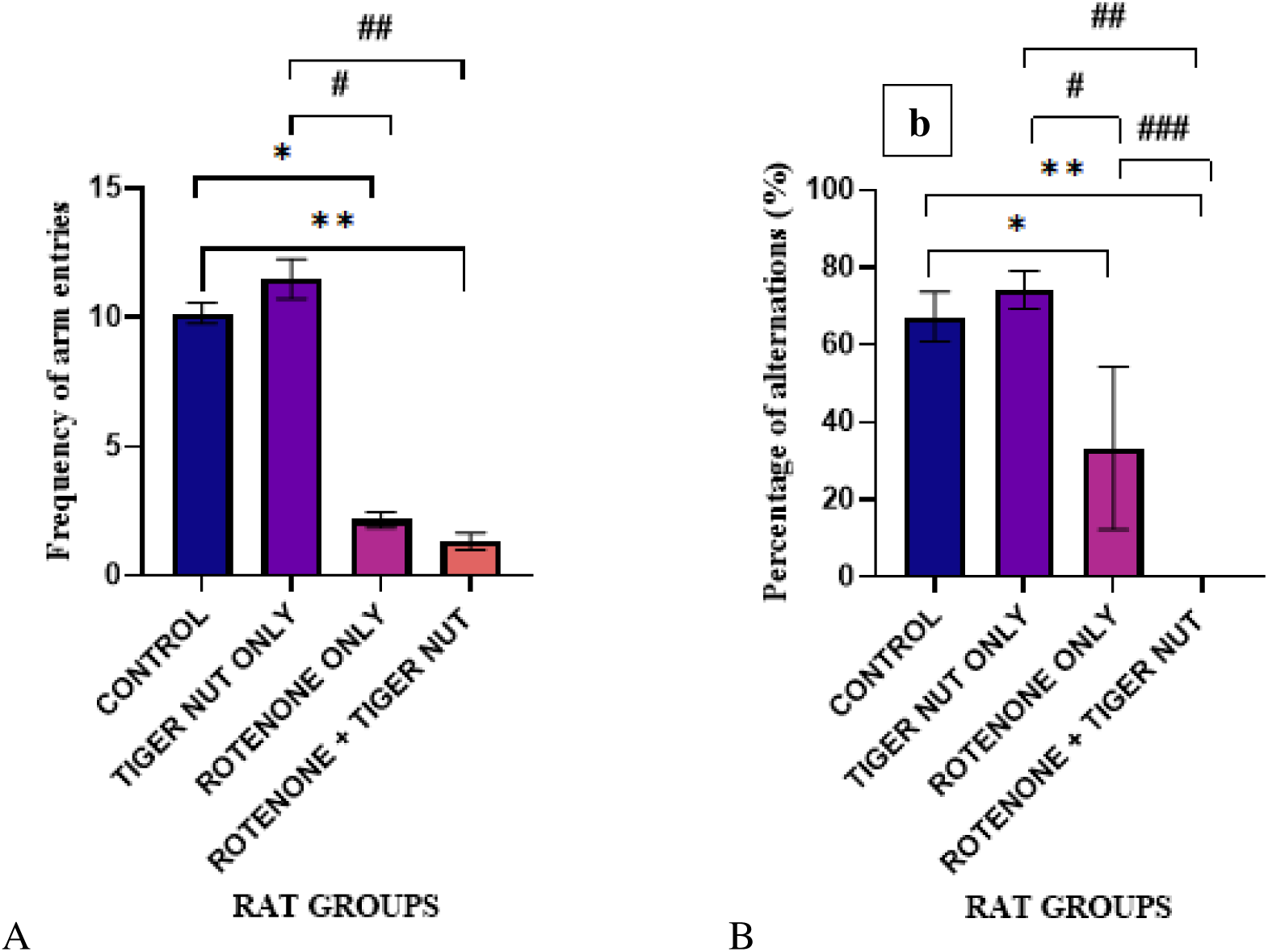
(a & b): Bar charts showing (a) the total arm entries and (b) the percentage of alternations (%) of the rats across experimental groups. Values are presented as Mean ± SEM. P ≤ 0.05.

#### 3.2.2 Open Field Test

Figure 3a and 3b show the rearing frequency and number of lines crossed by the rats across the experimental groups respectively. The results showed a significant increase in rearing in the control and rotenone only groups when compared with the rotenone plus tiger nut group, while the rotenone plus tiger nut group also showed a significant reduction when compared with the tiger nut only group. Regarding the number of lines crossed, the results showed a significant increase in the control and tiger nut only groups when compared with both the rotenone only and rotenone plus tiger nut groups.

**Figure 3.**
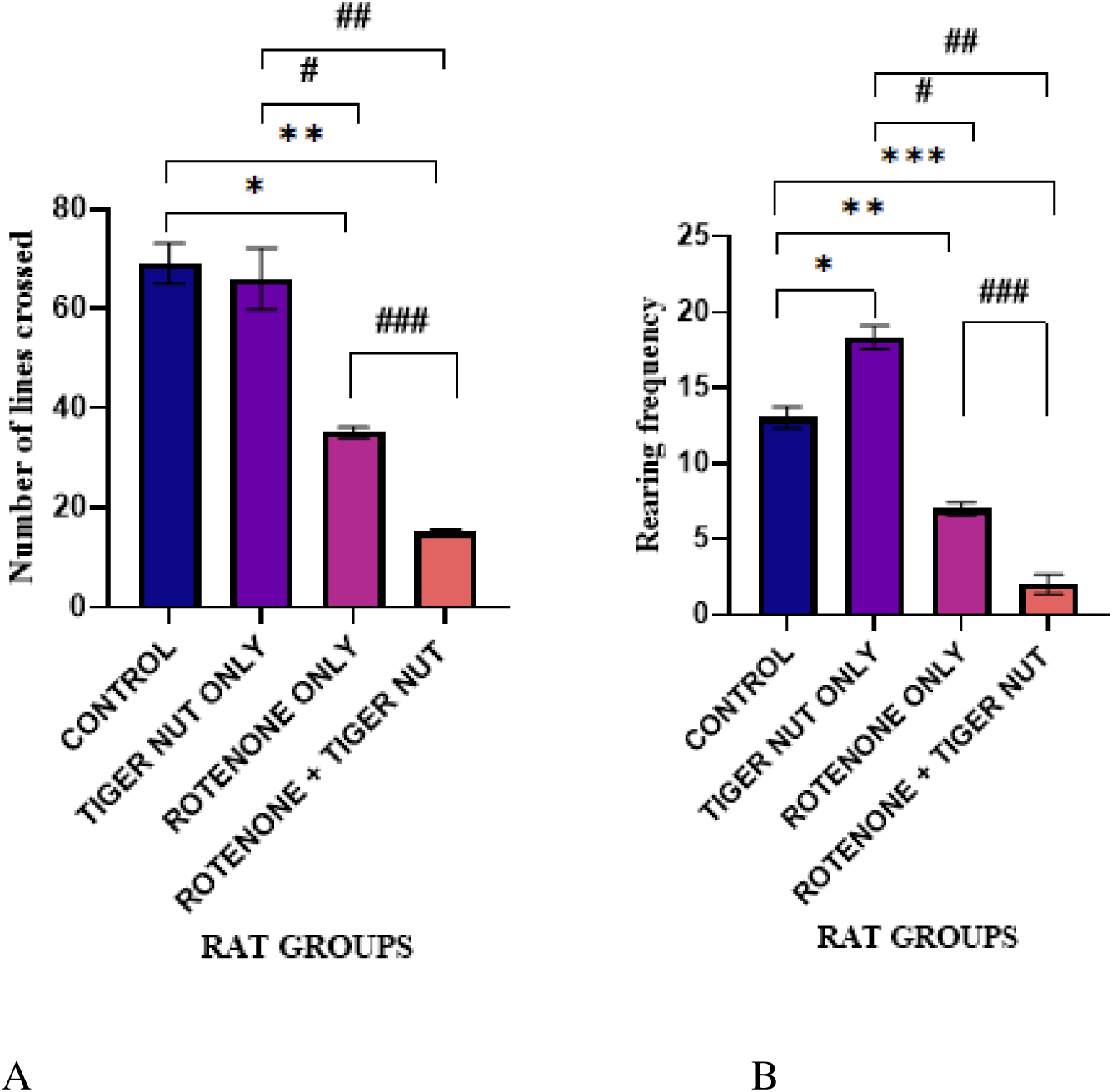
(a & b): Bar charts showing (a) the rearing frequency and (b) the number of lines crossed by the rats across experimental groups. Values are presented as Mean ± SEM. P ≤ 0.05.

### 3.3 Biochemical Assay Results

Figure 4 shows the catalase (CAT) activity across all experimental groups. The results showed a significant increase in the control group when compared with the rotenone only group. The tiger nut only group showed a significant reduction when compared with the rotenone plus tiger nut group. The rotenone only group also showed a significant reduction when compared with the rotenone plus tiger nut group, which recorded the highest catalase activity across all groups.

**Figure 4:**
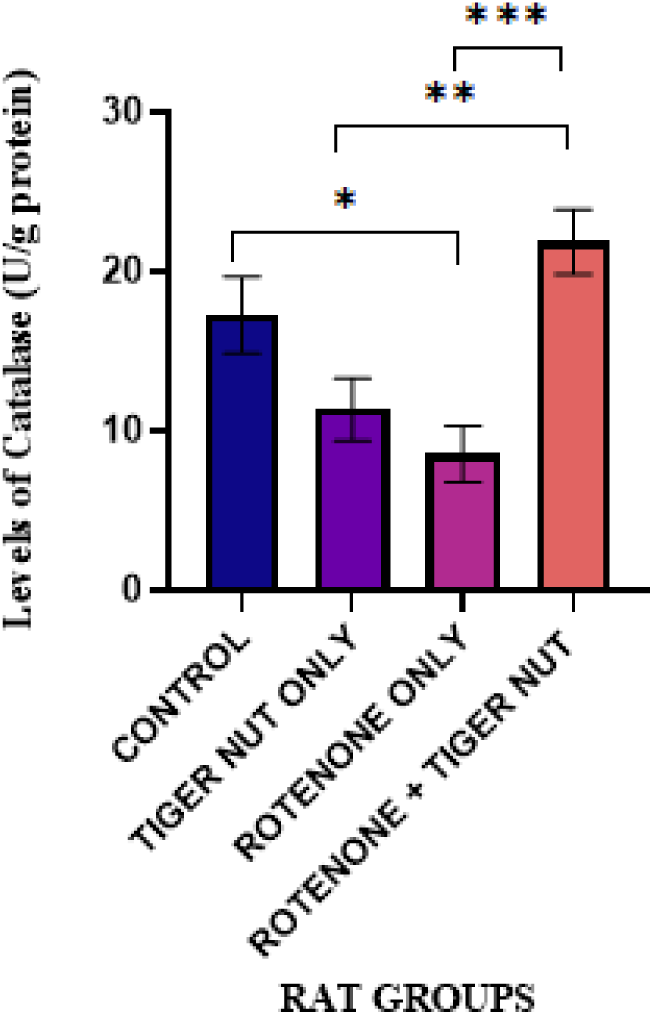
Bar chart showing the catalase (CAT) activity across all experimental groups. Values are expressed as Mean ± SEM. P ≤ 0.05.

### 3.4 Neurotransmitter Assay Results

Figure 5 shows the levels of gamma-aminobutyric acid (GABA) across all experimental groups. The results showed no significant difference across all groups. However, an observable increase was noted in the control group compared to the tiger nut only, rotenone only, and rotenone plus tiger nut groups.

**Figure 5:**
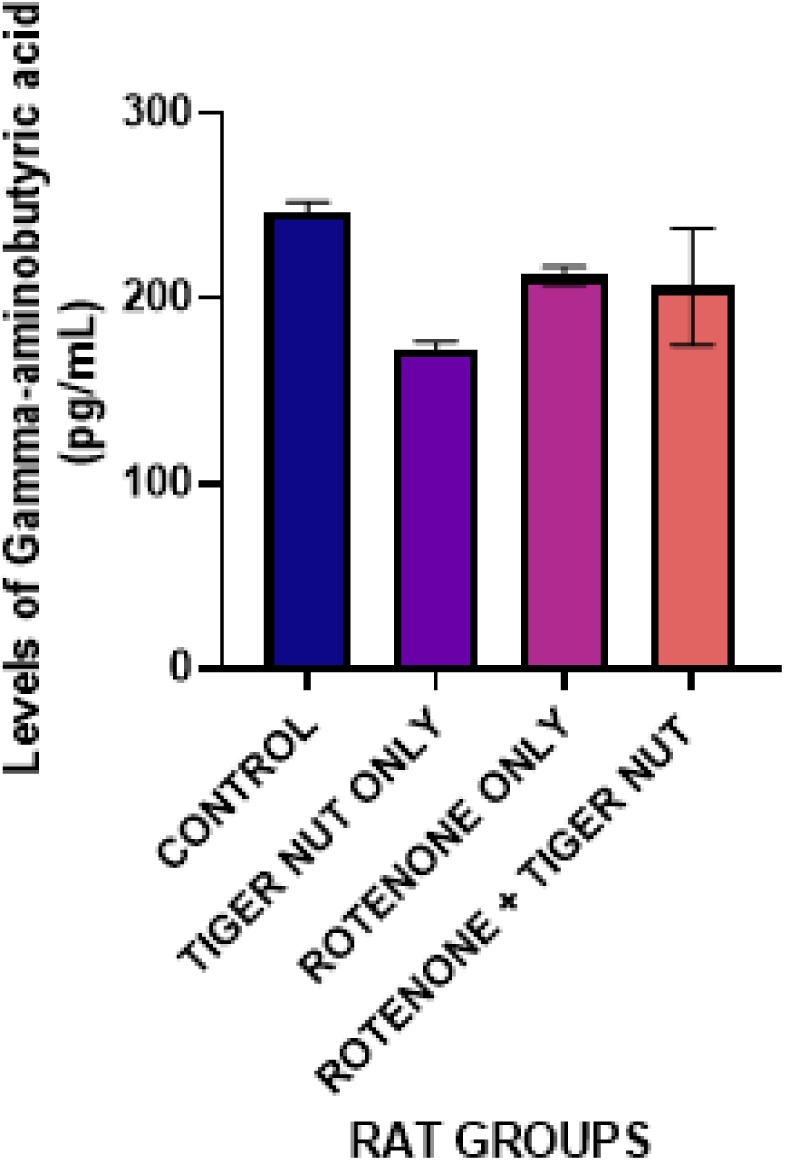
Bar chart showing the levels of gamma-aminobutyric acid (GABA) across all experimental groups. Values are expressed as Mean ± SEM. There was no significant difference observed. P ≤ 0.05.

### 3.5 Histological Analysis

#### 3.5.1 Hematoxylin and Eosin (H&E)

Plates 1 and 2 show the panoramic and high magnification views of H&E-stained hippocampal sections across the experimental groups respectively. The results showed that the control and tiger nut only groups displayed intact pyramidal neurons with well-defined nuclei and densely compacted cells across the CA1, CA2, CA3, and dentate gyrus subfields. The rotenone only group showed prominent vacuolation of pyramidal neurons, particularly in the CA1 and CA3 subfields. The rotenone plus tiger nut group showed near-normal histoarchitecture with a notable reduction in vacuolation compared to the rotenone only group.

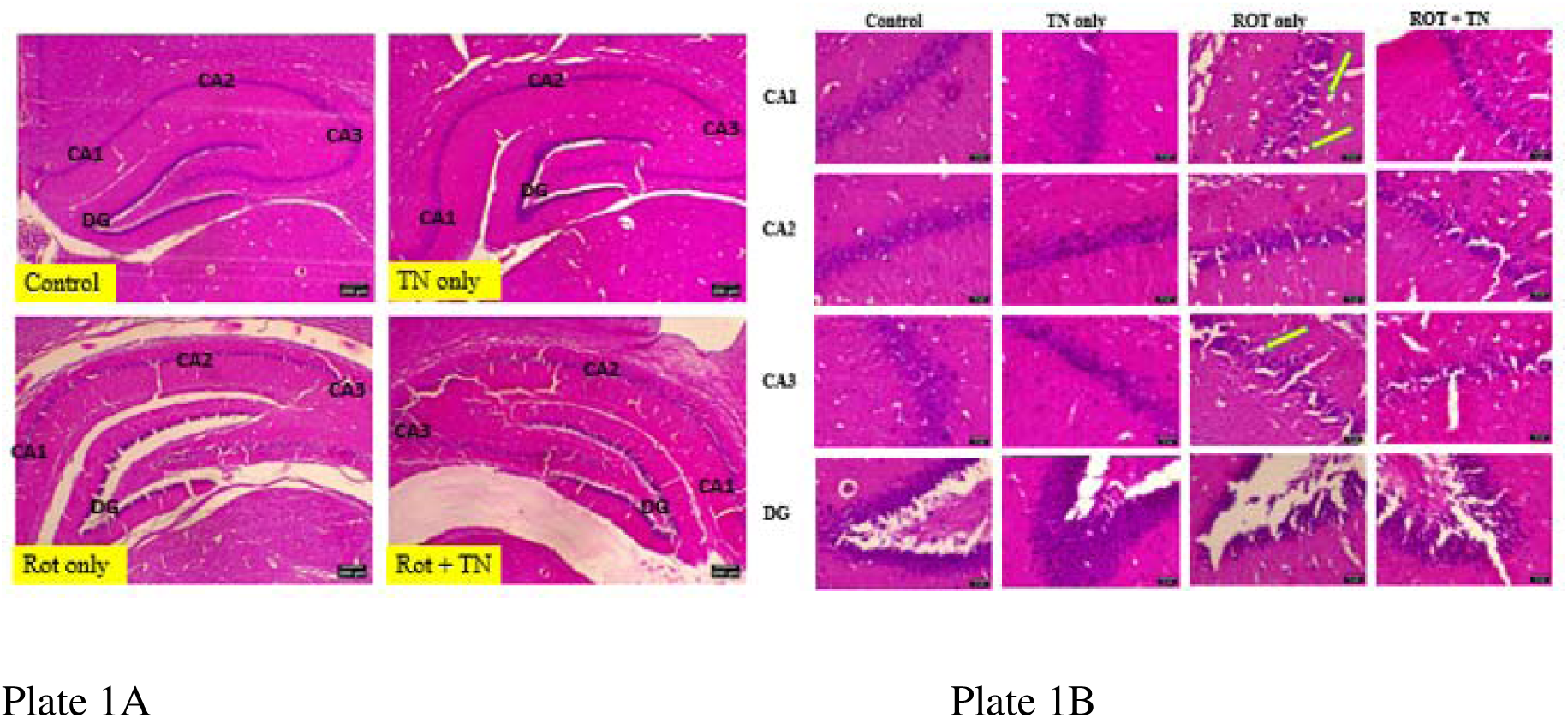

Plate 1A: Photomicrograph of H&E-stained hippocampal sections showing the panoramic view of the Cornu Ammonis (CA1, CA2, and CA3) and dentate gyrus across experimental groups. TN = tiger nut; ROT = rotenone. Scale bar = 200 µm.

Plate 1B: Photomicrograph of H&E-stained hippocampal sections showing pyramidal neuronal cells in the CA1–CA3 and dentate gyrus. Cells in the CA1 and CA3 of the rotenone only group show vacuolation (arrows). The rotenone plus tiger nut group shows near-normal histoarchitecture. TN = tiger nut; ROT = rotenone. Scale bar = 32 µm

#### 3.5.2 Cresyl Fast Violet (CFV)

Plates 3 and 4 show the panoramic and high magnification views of CFV-stained hippocampal sections across the experimental groups respectively. The results showed that the control and tiger nut only groups displayed pyramidal neurons with intact and well-defined Nissl bodies across all hippocampal subfields. The rotenone only group showed a notable loss of Nissl substance in the CA1, CA2, and CA3 subfields, while the dentate gyrus remained intact and comparable to the control. The rotenone plus tiger nut group showed some loss of Nissl substance in the CA1, CA2, and CA3 subfields, though the dentate gyrus remained intact.

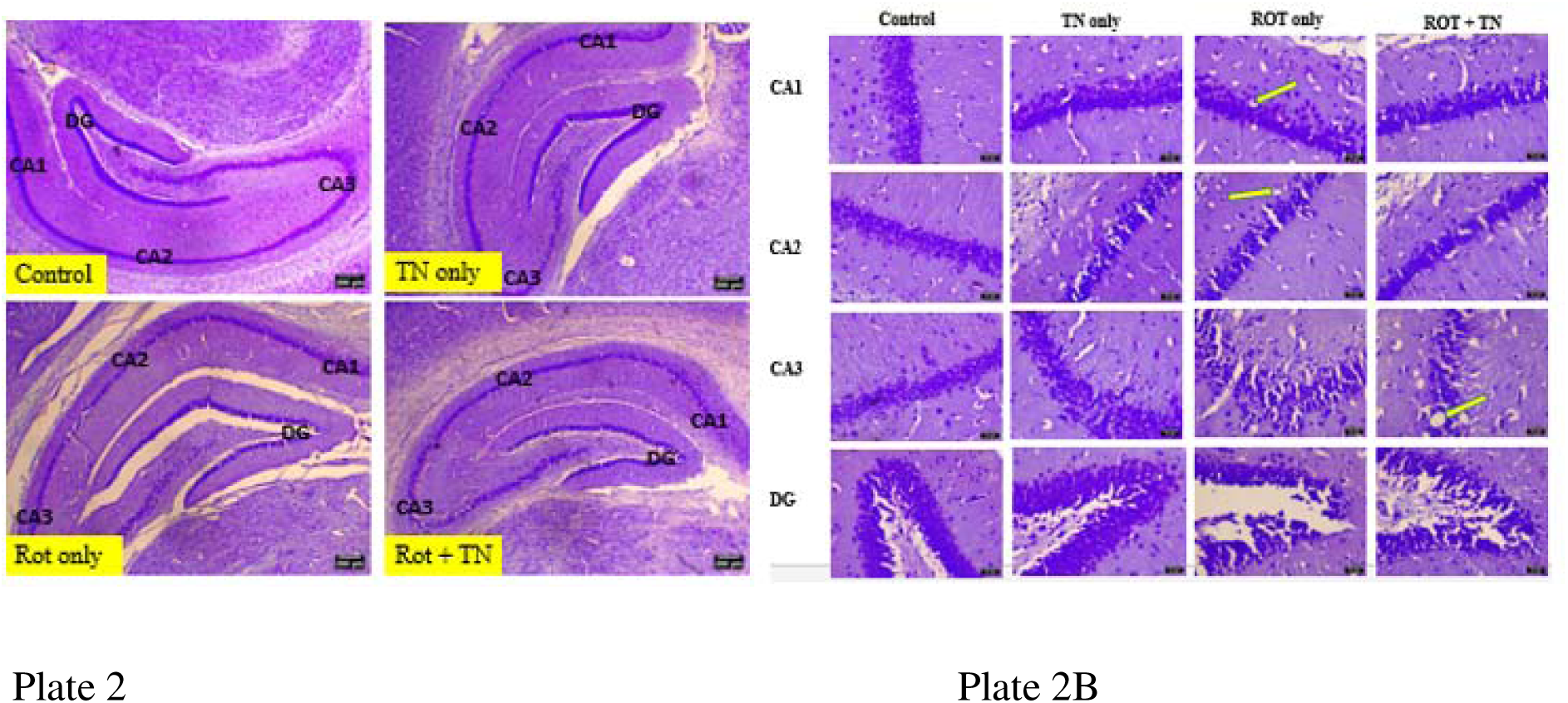

Plate 3: Photomicrograph of CFV-stained hippocampal sections showing the panoramic view of the Cornu Ammonis (CA1, CA2, and CA3) and dentate gyrus across experimental groups. TN = tiger nut; ROT = rotenone. Scale bar = 200 µm.

Plate 4: Photomicrograph of CFV-stained hippocampal sections showing Nissl substance distribution. The rotenone only and rotenone plus tiger nut groups show loss of Nissl bodies in the CA1, CA2, and CA3 subfields (arrows), with the dentate gyrus remaining intact. TN = tiger nut; ROT = rotenone. Scale bar = 32 µm.

#### 3.5.3 Luxol Fast Blue (LFB)

Plates 5 and 6 show the panoramic and high magnification views of LFB-stained hippocampal sections across the experimental groups respectively. The results showed that the control and tiger nut only groups demonstrated well-delineated myelin staining across all hippocampal subfields. The rotenone only group showed reduced myelin staining and disrupted hippocampal architecture compared to the control. The rotenone plus tiger nut group showed relatively well-delineated hippocampal architecture with appreciable myelin staining, indicating partial myelin preservation compared to the rotenone only group.

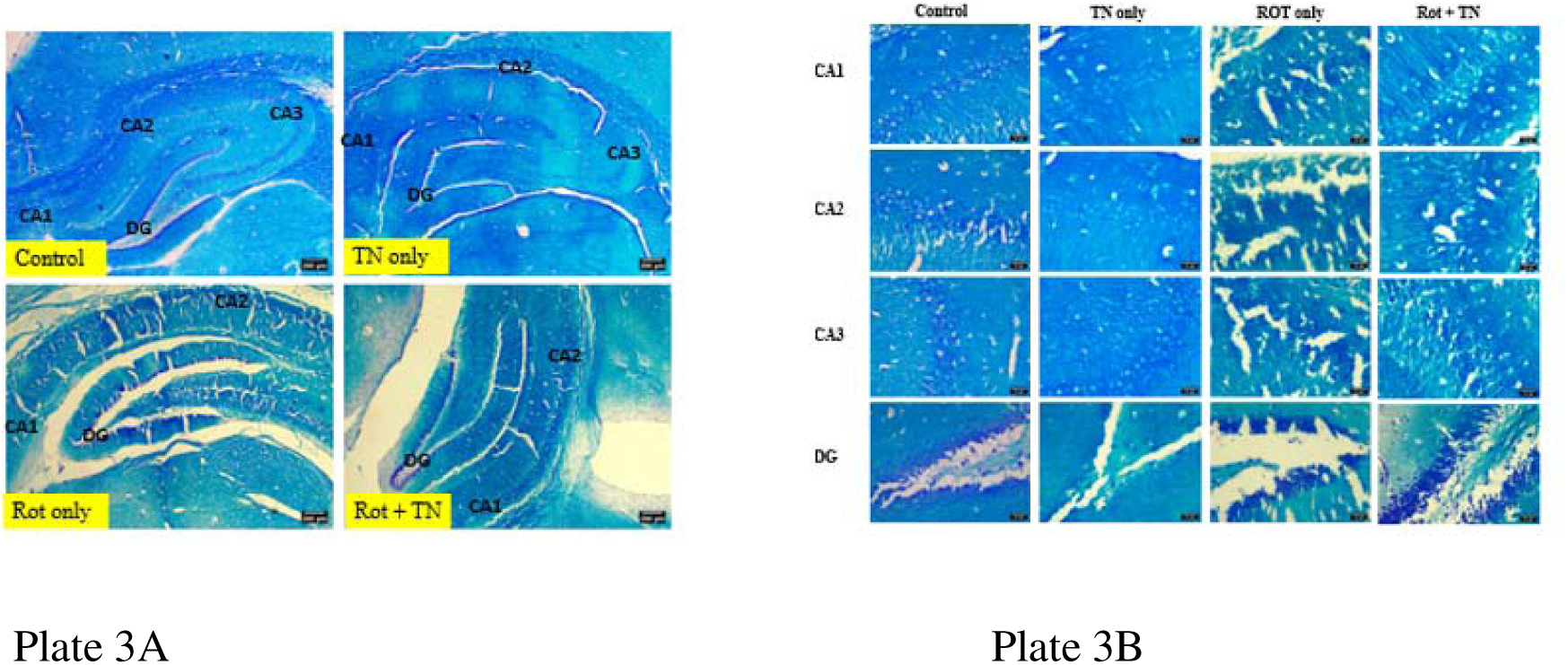

Plate 3A: Photomicrograph of LFB-stained hippocampal sections showing the panoramic view of the Cornu Ammonis (CA1, CA2, and CA3) and dentate gyrus across experimental groups. TN = tiger nut; ROT = rotenone. Scale bar = 200 µm.

Plate 3B: Photomicrograph of LFB-stained hippocampal sections showing myelin sheath distribution across experimental groups. TN = tiger nut; ROT = rotenone. Scale bar = 32 µm.

## Discussion

This study investigated the neuroprotective efficacy of tiger nut (*Cyperus esculentus*)-infused diet on the hippocampus in a rotenone-induced Parkinson’s disease model using male Wistar rats, evaluating morphological, neurobehavioral, biochemical, neurotransmitter, and histological outcomes. The results show that rotenone administration produced pronounced neurotoxic effects across all assessed parameters, while tiger nut-infused diet offered partial neuroprotection, most consistently evidenced by enhanced catalase activity and improved hippocampal cytoarchitecture in the rotenone plus tiger nut group.

The significant reduction in body weight observed in the rotenone only group is consistent with rotenone’s ability to disrupt cellular energy metabolism and impair gastrointestinal function, leading to reduced appetite and progressive weight loss in treated animals, as previously reported [20]. The rotenone plus tiger nut group recorded the lowest body weight change of all groups, suggesting that tiger nut did not attenuate rotenone-induced weight loss and may have exacerbated it. This contrasts with findings by [21], who demonstrated that dietary protein supplementation reversed rotenone-induced weight reduction, underscoring that the nutritional and neuroprotective benefits of dietary interventions are highly dependent on the nature and composition of the supplement used. The unexpectedly lower body weight in the tiger nut only group compared to the control may reflect differences in bioavailability and caloric density between a tiger nut-infused diet and the tiger nut oil-based preparations used in earlier studies that reported weight gain [22], and warrants further investigation.

The increased brain weight in the rotenone only and rotenone plus tiger nut groups is consistent with rotenone-induced neuroinflammation, which drives microglial activation, reactive astrogliosis, and alpha-synuclein aggregation, collectively increasing brain tissue volume [23]. The absence of a significant difference between these two groups suggests that tiger nut supplementation did not significantly attenuate the neuroinflammatory mass changes at the dose and duration used. The lowest brain weight recorded in the tiger nut only group may reflect a modulatory influence of tiger nut on brain lipid composition under normal physiological conditions, given the high oleic acid content of Cyperus esculentus [15].

The significantly reduced arm entries, alternations, and percentage of alternations in the rotenone only group confirm that rotenone produced both motor and cognitive impairment, consistent with reports that rotenone-induced dopaminergic neurodegeneration causes bradykinesia and hippocampal cognitive dysfunction [8, 24]. The tiger nut only group maintained normal Y-maze performance, in agreement with findings that tiger nut extract restored memory impairment and reduced acetyl cholinesterase activity in the hippocampus of rats [17, 25]. The rotenone plus tiger nut group failed to show significant recovery across all Y-maze parameters, suggesting that the severity of rotenone-induced neurodegeneration in this model may require a higher dose or longer duration of supplementation to achieve meaningful cognitive restoration [16]. The lower alternate arm return in the rotenone-induced groups further reflects the disruption of spatial memory and environmental cue utilization consistent with hippocampal oxidative damage [26].

The significantly reduced rearing and total locomotor activity in the rotenone only and rotenone plus tiger nut groups in the open field test are consistent with well-established reports that rotenone reduces spontaneous motor activity through dopaminergic cell loss and mitochondrial dysfunction [24, 27]. The normal locomotor performance in the tiger nut only group supports the view that tiger nut does not impair motor function and may even support it through its antioxidant and neuroprotective constituents [15]. The reduced defecation in the rotenone only group may reflect rotenone-induced autonomic nervous system dysfunction, while the lower defecation in the tiger nut only group may indicate a calming modulatory effect on autonomic reactivity [24, 15]. The higher defecation in the rotenone plus tiger nut group compared to both the rotenone only and tiger nut only groups may reflect a heightened autonomic response arising from an interaction between rotenone-induced neurodegeneration and tiger nut supplementation that warrants further investigation.

Although MDA levels and GSH showed no statistically significant differences across groups, the observable decrease in MDA in the rotenone plus tiger nut group and the prevention of further GSH depletion in this group suggest that tiger nut-infused diet provided a modest degree of antioxidant protection against rotenone-induced oxidative damage. These trends are consistent with reports that tiger nut administration reduced lipid peroxidation products and activated the Keap1-Nrf2 pathway to support glutathione metabolism in animals subjected to toxic insults [31]. The significant reduction in catalase activity in the rotenone only group reflects rotenone-induced suppression of antioxidant enzyme function through excessive reactive oxygen species generation, as previously reported [29, 30]. Notably, the rotenone plus tiger nut group recorded the highest catalase activity of all groups, suggesting that the combination of rotenone-induced oxidative stress and tiger nut supplementation stimulated a compensatory upregulation of catalase activity, consistent with reports that Cyperus esculentus extract significantly increased catalase and other antioxidant enzyme activities in animals exposed to toxic insults [31, 32]. The absence of significant differences in GSH levels, alongside the observable reduction in the rotenone-induced groups, is consistent with rotenone’s established mechanism of GSH depletion through mitochondrial complex I inhibition [33], while the maintenance of GSH in the rotenone plus tiger nut group supports the Keap1-Nrf2 activating capacity of tiger nut constituents [31].

The observable reduction in GABA levels in the rotenone only and rotenone plus tiger nut groups reflects the disruption of inhibitory neurotransmission associated with rotenone-induced dopaminergic neurodegeneration, consistent with reports that GABAergic signaling abnormalities in the basal ganglia are closely associated with both motor and non-motor symptoms of Parkinson’s disease [28]. The slightly higher GABA in the rotenone plus tiger nut group compared to the rotenone only group, though not statistically significant, suggests a modest modulatory effect of tiger nut on GABAergic neurotransmission, consistent with findings that tiger nut extract restored neurotransmitter-related impairments in the context of neurotoxic challenge [25].

Histological examination confirmed the neurotoxic effects of rotenone at the structural level. The neuronal vacuolation observed in the CA1 and CA3 subfields on H&E staining, and the chromatolysis evidenced by Nissl substance loss on CFV staining in the rotenone only group, are consistent with rotenone-induced mitochondrial dysfunction and oxidative neuronal injury [34]. The relative preservation of the dentate gyrus across all staining techniques and groups suggests differential subfield vulnerability to rotenone-induced hippocampal damage, consistent with established patterns of hippocampal regional susceptibility [6]. In the rotenone plus tiger nut group, near-normal histoarchitecture on H&E staining, partial Nissl body preservation on CFV staining, and relatively well-maintained myelin distribution on LFB staining collectively indicate that tiger nut supplementation attenuated rotenone-induced hippocampal structural damage, likely through its antioxidant and anti-inflammatory constituents including flavonoids, phenolic acids, and vitamin E [35].

## Strengths of the Study

A major strength of this study is the use of a multidimensional experimental design that integrated morphological, neurobehavioral, biochemical, neurotransmitter, histological, and immunohistochemical assessments. This comprehensive approach allowed the neurotoxic effects of rotenone and the potential neuroprotective effects of tiger nut-infused diet to be evaluated simultaneously across multiple levels of biological organization, from systemic weight changes to hippocampal cytoarchitecture, lending greater interpretive depth and reliability to the findings.

The rotenone model employed in this study is one of the most widely validated and translatable experimental models of Parkinson’s disease, as it reliably reproduces both motor and non-motor features of the disease through mitochondrial complex I inhibition, oxidative stress, and progressive dopaminergic neurodegeneration [34]. The use of this model therefore strengthens the translational relevance of the findings.

The inclusion of tiger nut as a dietary intervention in its naturally consumed infused-diet form, rather than as an isolated extract, reflects ecologically valid and practically accessible conditions of administration. This approach is directly relevant to the assessment of tiger nut as a complementary dietary strategy in real-world contexts, where populations consume it as a food source rather than a pharmaceutical preparation [15].

The selection of hippocampal sub regions for analysis, including CA1, CA2, CA3, and the dentate gyrus, represents another notable strength, as it allowed for the characterization of differential subfield vulnerability to rotenone-induced neurotoxicity and the regional neuroprotective capacity of tiger nut supplementation, consistent with established functional and structural heterogeneity across hippocampal subfields [6].

Finally, the assessment of parvalbumin immunoexpression provided an additional layer of sensitivity in evaluating interneuronal integrity, as parvalbumin-expressing GABAergic interneurons are among the most vulnerable neuronal populations in neurodegenerative conditions and represent a functionally critical component of hippocampal inhibitory circuitry [36].

## Conclusion

This study investigated the neuroprotective efficacy of tiger nut (*Cyperus esculentus*)-infused diet on the hippocampus in a rotenone-induced Parkinson’s disease model using male Wistar rats. The findings demonstrate that chronic rotenone administration successfully induced Parkinson’s disease-like features in the experimental animals, as evidenced by significant impairments in locomotor activity and spatial working memory, significant reduction in catalase activity, and marked histopathological alterations in the hippocampus including neuronal vacuolation, chromatolysis, and myelin disruption, particularly in the CA1 and CA3 subfields [34].

Tiger nut-infused diet demonstrated partial neuroprotective potential against rotenone-induced hippocampal neurotoxicity. This was most notably evidenced by the significant enhancement of catalase activity and the improved preservation of hippocampal cytoarchitecture observed in the rotenone plus tiger nut group compared to the rotenone only group [31, 32]. However, full restoration of all behavioral, biochemical, neurotransmitter, and histological parameters to control levels was not achieved, and the neuroprotection is therefore classified as partial rather than complete.

These findings suggest that tiger nut-infused diet, through its rich content of antioxidant and anti-inflammatory bioactive compounds, can partially attenuate the oxidative stress and structural damage inflicted on the hippocampus by rotenone, and may therefore hold promise as a complementary dietary intervention in the management of Parkinson’s disease-related cognitive decline and hippocampal dysfunction [15, 35]. Further studies focusing on dose optimization, molecular mechanisms, and validation in higher experimental models are recommended to fully explore the neuroprotective capacity of tiger nut.

## References

1. Poewe, W., Seppi, K., Tanner, C. M., Halliday, G. M., Brundin, P., Volkmann, J., & Lang, A. E. (2017). Parkinson disease. Nature Reviews Disease Primers, 3(1), 17013. 10.1038/nrdp.2017.13

2. Aarsland, D., Batzu, L., Halliday, G. M., Geurtsen, G. J., Ballard, C., Ray Chaudhuri, K., & Weintraub, D. (2021). Parkinson disease-associated cognitive impairment. Nature Reviews Disease Primers, 7(1), 47. 10.1038/s41572-021-00280-3

3. Dorsey, E. R., & Bloem, B. R. (2018). The Parkinson pandemic — a call to action. JAMA Neurology, 75(1), 9–10. 10.1001/jamaneurol.2017.3299

4. Feigin, V. L., Nichols, E., Alam, T., Bannick, M. S., Beghi, E., Blake, N., & Fischer, F. (2019). Global, regional, and national burden of neurological disorders, 1990–2016: A systematic analysis for the Global Burden of Disease Study 2016. The Lancet Neurology, 18(5), 459–480. 10.1016/S1474-4422(18)30499-X

5. Dorsey, E. R., Elbaz, A., Nichols, E., Abd-Allah, F., Abdelalim, A., Adsuar, J. C., & Murray, C. J. (2018). Global, regional, and national burden of Parkinson’s disease, 1990–2016: A systematic analysis for the Global Burden of Disease Study 2016. The Lancet Neurology, 17(11), 939–953. 10.1016/S1474-4422(18)30295-3

6. Strange, B. A., Witter, M. P., Lein, E. S., & Moser, E. I. (2014). Functional organization of the hippocampal longitudinal axis. Nature Reviews Neuroscience, 15(10), 655–669. 10.1038/nrn3785

7. Dickson, D. W. (2018). Neuropathology of Parkinson disease. Parkinsonism & Related Disorders, 46, S30–S33. 10.1016/j.parkreldis.2017.07.033

8. Guo, Z., Ruan, Z., Zhang, D., Liu, X., Hou, L & Wang, Q. (2022). Rotenone impairs learning and memory in mice through microglia-mediated blood brain barrier disruption and neuronal apoptosis. Chemosphere, 291, 132982. 10.1016/j.chemosphere.2021.132982

9. Darbinyan, L. V., Hambardzumyan, L. E., Simonyan, K. V., Chavushyan, V. A., Manukyan, L. P., & Sarkisian, V. H. (2017). Rotenone impairs hippocampal neuronal activity in a rat model of Parkinson’s disease. Pathophysiology, 24(1), 23–30. 10.1016/j.pathophys.2016.12.001

10. Innos, J., & Hickey, M. A. (2021). Using rotenone to model Parkinson’s disease in mice: A review of the role of pharmacokinetics. Chemical Research in Toxicology, 34(5), 1223–1239. 10.1021/acs.chemrestox.0c00522

11. Ascherio, A., & Schwarzschild, M. A. (2016). The epidemiology of Parkinson’s disease: Risk factors and prevention. The Lancet Neurology, 15(12), 1257–1272. 10.1016/S1474-4422(16)30230-7

12. Connolly, B. S., & Lang, A. E. (2014). Pharmacological treatment of Parkinson disease: A review. JAMA, 311(16), 1670–1683. 10.1001/jama.2014.3654

13. Arafat, S. M., Gaafar, A. M., Basuny, A. M., & Nassef, S. L. (2009). Chufa tubers (Cyperus esculentus L.): As a new source of food. World Applied Sciences Journal, 7(2), 151–156.

14. Sánchez-Zapata, E., Fernández-López, J., & Angel Pérez-Álvarez, J. (2012). Tiger nut (Cyperus esculentus) commercialization: Health aspects, composition, properties, and food applications. Comprehensive Reviews in Food Science and Food Safety, 11(4), 366–377. 10.1111/j.1541-4337.2012.00190.x

15. Edo, G. I., Samuel, P. O., Nwachukwu, S. C., Ikpekoro, V. O., Promise, O., Oghenegueke, O., & Ajakaye, R. S. (2024). A review on the biological and bioactive components of Cyperus esculentus L.: Insight on food, health, and nutrition. Journal of the Science of Food and Agriculture, 104(14), 8414–8429. 10.1002/jsfa.13666

16. Umukoro, S., Okoh, L., Igweze, S. C., Ajayi, A. M., & Ben-Azu, B. (2020). Protective effect of Cyperus esculentus (tiger nut) extract against scopolamine-induced memory loss and oxidative stress in mouse brain. Drug Metabolism and Personalized Therapy, 35(3), 20200112. 10.1515/dmpt-2020-0112

17. Arogundade, T., Yawson, E., Gbadamosi, I., Balogun, W., Akinola, O., Obasi, K., & Lambe, E. (2018). Behavioural cellular and neurochemical alterations in rat prefrontal cortex and hippocampus exposed to tigernut (Cyperus esculentus) treatment. Journal of Environmental Toxicology and Public Health, 3, 38–46.

18. Olabiyi, A. A., Oboh, G., Akinyemi, A. J., Ademiluyi, A. O., Boligon, A. A., & de Campos, M. M. A. (2017). Tiger nut (Cyperus esculentus L.) supplemented diet modulate key biochemical indices relevant to erectile function in male rats. Journal of Functional Foods, 34, 152–158. 10.1016/j.jff.2017.04.026

19. Swamy, G., Holla, R., & Rao, S. R. (2021). Establishing the rotenone-induced Parkinson’s disease animal model in Wistar Albino rats. Journal of Health and Allied Sciences NU, 11(03), 158–163. 10.1055/s-0041-1726125

20. Dos Santos, J. C. C., Reboucas, C. D. S. M., Oliveira, L. F., dos Santos Cardoso, F., de Souza Nascimento, T., Oliveira, A. V., & de Barros Viana, G. S. (2023). The role of the gut-brain axis in a rotenone-induced rat model of Parkinson’s disease. Neurobiology of Aging, 132, 185–197. 10.1016/j.neurobiolaging.2023.08.009

21. Zarei, P., Amirpour-Najafabadi, B., Sam-Sani, P., Sakhaie, M. H., & Sadegh, M. (2024). Protective effect of whey protein supplement against rotenone-induced motor dysfunction in a rat model of Parkinson’s disease. Advanced Biomedical Research, 13(1), 93. 10.4103/abr.abr_23_23

22. Bamishaiye, E. I., Muhammad, N. O., & Bamishaiye, O. M. (2010). Assessment of the biological value of tiger nut (Cyperus esculentus) tuber oil meal-based diet in rats. Annals of Biological Research, 1(4), 274–280.

23. Rocha, S. M., Bantle, C. M., Aboellail, T., Chatterjee, D., Smeyne, R. J., & Tjalkens, R. B. (2022). Rotenone induces regionally distinct α-synuclein protein aggregation and activation of glia prior to loss of dopaminergic neurons in C57Bl/6 mice. Neurobiology of Disease, 167, 105685. 10.1016/j.nbd.2022.105685

24. Sheta, R., Bérard, M., Musiol, D., Martínez-Drudis, L., & Oueslati, A. (2024). Behavioral analysis of motor and non-motor impairment in rodent models of Parkinson’s disease. Frontiers in Aging Neuroscience, 16, 1464706. 10.3389/fnagi.2024.1464706

25. Saeed, M. M., Fernández-Ochoa, Á., Saber, F. R., Sayed, R. H., Cádiz-Gurrea, M. L., Elmotayam, A. K., Leyva-Jiménez, F. J., Segura-Carretero, A., & Nadeem, R. I. (2022). The potential neuroprotective effect of Cyperus esculentus L. extract in scopolamine-induced cognitive impairment in rats: Extensive biological and metabolomics approaches. Molecules, 27(20), 7118. 10.3390/molecules27207118

26. Vastegani, S. M., Khoshnam, S. E., Mansouri, E., Hajipour, S., Ghafouri, S., Bakhtiari, N., & Farbood, Y. (2023). Neuroprotective effect of anethole against rotenone-induced non-motor deficits and oxidative stress in rat model of Parkinson’s disease. Behavioural Brain Research, 437, 114100. 10.1016/j.bbr.2022.114100

27. Sun, C., Wang, Y., Mo, M., Song, C., Wang, X., Chen, S., & Liu, Y. (2019). Minocycline protects against rotenone-induced neurotoxicity correlating with upregulation of Nurr1 in a Parkinson’s disease rat model. BioMed Research International, 2019, 6843265. 10.1155/2019/6843265

28. Alharbi, B., Al-Kuraishy, H. M., Al-Gareeb, A. I., Elekhnawy, E., Alharbi, H., Alexiou, A., & Batiha, G. E. S. (2024). Role of GABA pathway in motor and non-motor symptoms in Parkinson’s disease: A bidirectional circuit. European Journal of Medical Research, 29(1), 205. 10.1186/s40001-024-01798-y

29. Kryl’skii, E. D., Razuvaev, G. A., Potapova, T. N., Akinina, A. I., & Nihaev, L. E. (2021). Functioning of the antioxidant defense system in rotenone-induced Parkinson’s disease. Bulletin of Experimental Biology and Medicine, 171(6), 716–721. 10.1007/s10517-021-05290-3

30. Singh, R., Zahra, W., Singh, S. S., Birla, H., Rathore, A. S., Keshri, P. K., & Singh, S. P. (2023). Oleuropein confers neuroprotection against rotenone-induced model of Parkinson’s disease via BDNF/CREB/Akt pathway. Scientific Reports, 13(1), 2452. 10.1038/s41598-023-29287-4

31. He, L., Tan, W., Wei, C., Dai, F., & Tian, J. (2025). Tiger nut (Cyperus esculentus L.): A systematic review of metabolic regulation, functional bioactivities, and sustainable food applications. Journal of Food Science, 90(11), e70705. 10.1111/1750-3841.70705

32. Udefa, A. L., Amama, E. A., Archibong, E. A., Nwangwa, J. N., Adama, S., Inyang, V. U., & Inah, I. O. (2020). Antioxidant, anti-inflammatory and anti-apoptotic effects of hydro-ethanolic extract of Cyperus esculentus L. (tigernut) on lead acetate-induced testicular dysfunction in Wistar rats. Biomedicine & Pharmacotherapy, 129, 110491. 10.1016/j.biopha.2020.110491

33. Yulianti, A. B., Sumarsono, S. H., Ridwan, A., & Yusuf, A. T. (2021). Protofibril formation: Decreased total glutathione concentration as an early indicator of neuron damage in the brainstems of Wistar rats treated with rotenone. F1000Research, 10, 1158. 10.12688/f1000research.54449.2

34. Cannon, J. R., Tapias, V., Na, H. M., Honick, A. S., Drolet, R. E., & Greenamyre, J. T. (2009). A highly reproducible rotenone model of Parkinson’s disease. Neurobiology of Disease, 34(2), 279–290. 10.1016/j.nbd.2009.01.016

35. Adejuyitan, J. A. (2011). Tigernut processing: Its food uses and health benefits. American Journal of Food Technology, 6(3), 197–201. 10.3923/ajft.2011.197.201

36. Hu, H., Gan, J., & Jonas, P. (2014). Fast-spiking, parvalbumin+ GABAergic interneurons: From cellular design to microcircuit function. Science, 345(6196), 1255263. 10.1126/science.1255263

